# Multi-level Analysis of Codon Usage Patterns Reveals Systematic Optimization of Oncogenic Gene Expression in Pancreatic Cancer

**DOI:** 10.64898/2026.04.24.720399

**Authors:** Lisa Müller, Markus Glaß, Pit Preckwinkel, Stefan Hüttelmaier, Monika Hämmerle, Tony Gutschner

**Affiliations:** Institute of Molecular Medicine, Section for RNA biology and pathogenesis, Faculty of Medicine, Martin Luther University Halle-Wittenberg, 06120 Halle, Germany; Institute of Molecular Medicine, Section for Molecular Cell Biology, Faculty of Medicine, Martin Luther University Halle-Wittenberg, 06120 Halle, Germany; Institute of Pathology, Section for Experimental Pathology, Faculty of Medicine, Martin Luther University Halle-Wittenberg, 06120 Halle, Germany

**Keywords:** Codon usage, codon bias, codon optimization, GC content, pancreatic cancer, PDAC, single-cell transcriptomics, tRNA, translation

## Abstract

**Background:** Codon usage bias, the non-random usage of synonymous codons in coding sequences, represents a fundamental feature of genomic organization that has been largely understudied in cancer biology. Pancreatic ductal adenocarcinoma (PDAC), the predominant subtype of pancreatic cancer, is characterized by aggressive disease progression and limited therapeutic options, necessitating novel approaches to understand its molecular pathogenesis. Leveraging publicly available single-cell RNA sequencing data, we performed comprehensive codon usage analyses across different cellular populations in PDAC.

**Results:** Employing a variety of computational codon usage indices uncovered the connections between cancer-specific cellular state features and codon usage signatures. Our findings reveal that malignant pancreatic cells express genes with significantly higher GC content, demonstrate preferential usage of optimal codons through increased frequency of preferred synonymous codons, and exhibit a marked preference for more cost-effective amino acids. Analysis of transcript-level bulk RNA-seq data from PDAC tumors revealed that these codon optimization patterns extend to alternative isoform usage, with highly expressed isoforms displaying increased codon optimality and enhanced mRNA stability.

**Conclusion:** These codon usage-dependent adaptations operating at both gene expression and transcript isoform levels may enable malignant cells to enhance gene expression rates, potentially leading to increased translational efficiency and protein production. These insights into the codon usage landscape of PDAC may provide potential biomarkers for disease monitoring and treatment response prediction.

## BACKGROUND

The phenomenon ‘codon usage bias’ describes the non-random or preferential use of synonymous codons (distinct codons that encode for the same amino acid) and is ubiquitous across genes and organisms. However, distinct species possess characteristic codon biases which are shaped by mutational pressure, natural selection, and random genetic drift (1, 2). Nonetheless, codon usage bias varies between genes within an organism, reflecting its role as a critical factor in determining gene expression by influencing mRNA stability, translation efficiency and accuracy, as well as the structure and hence functions of proteins (3). Whereas codon usage patterns at the genomic level across several taxa have already been investigated during the last decade (4), the influence of codon usage bias on pathological processes is not well understood.

Pancreatic ductal adenocarcinoma (PDAC) is the most prevalent neoplastic disease of the pancreas, and its global burden has increased dramatically over the past few decades (5). Despite existing studies on codon usage signatures in multiple tumor entities (6, 7), PDAC has not yet been included in pan-cancer analyses. We therefore investigated whether gene expression in PDAC might be influenced by codon usage patterns.

Dysregulated signal transduction and gene expression within primary tumor cells are critical for PDAC progression (8). Moreover, PDAC is characterized by high intra-tumoral heterogeneity (9). Since PDAC tumors are complex mixtures in which malignant epithelial cells often represent a minority of the tissue compartment but the stroma constitutes up to 70% of the tumor mass (10), this highly heterogeneous stroma poses challenges for identifying genetic signatures based on bulk mRNA sequencing. Thus, single-cell transcriptomics provides a powerful tool for exploring such complex systems, offering gene expression information at a single-cell resolution, and enabling researchers to distinguish different cell types and their functional states within tumor tissues. Peng *et al.* (11) employed a single-cell transcriptome approach to dissect PDAC intra-tumoral heterogeneity and associated critical factors of PDAC development.

To strengthen the understanding of the molecular mechanisms underlying malignancy in PDAC, we examined codon usage patterns within the gene expression landscape of PDAC by re-analyzing the Peng *et al.* data set (11) (**Figure 1**). Since ductal cells are considered one of the main cellular origins for PDAC formation (12, 13), we focused on two distinct populations: malignant and normal ductal cells. We employed various computational codon usage indices to uncover connections between cancer-specific cellular features and codon usage signatures. Our analysis revealed that malignant pancreatic cells exhibit distinct codon usage adaptations, including the preferential usage of optimal codons and more cost-effective amino acids. Complementary analysis of bulk RNA-seq data from PDAC tumors demonstrated that these optimization patterns extend to the transcript isoform level, where highly expressed isoforms display similar codon optimization features. These findings suggest that codon usage-dependent mechanisms, operating across multiple regulatory levels, may contribute to enhanced translational efficiency in PDAC and could provide potential biomarkers for disease monitoring.

**Figure 1:**
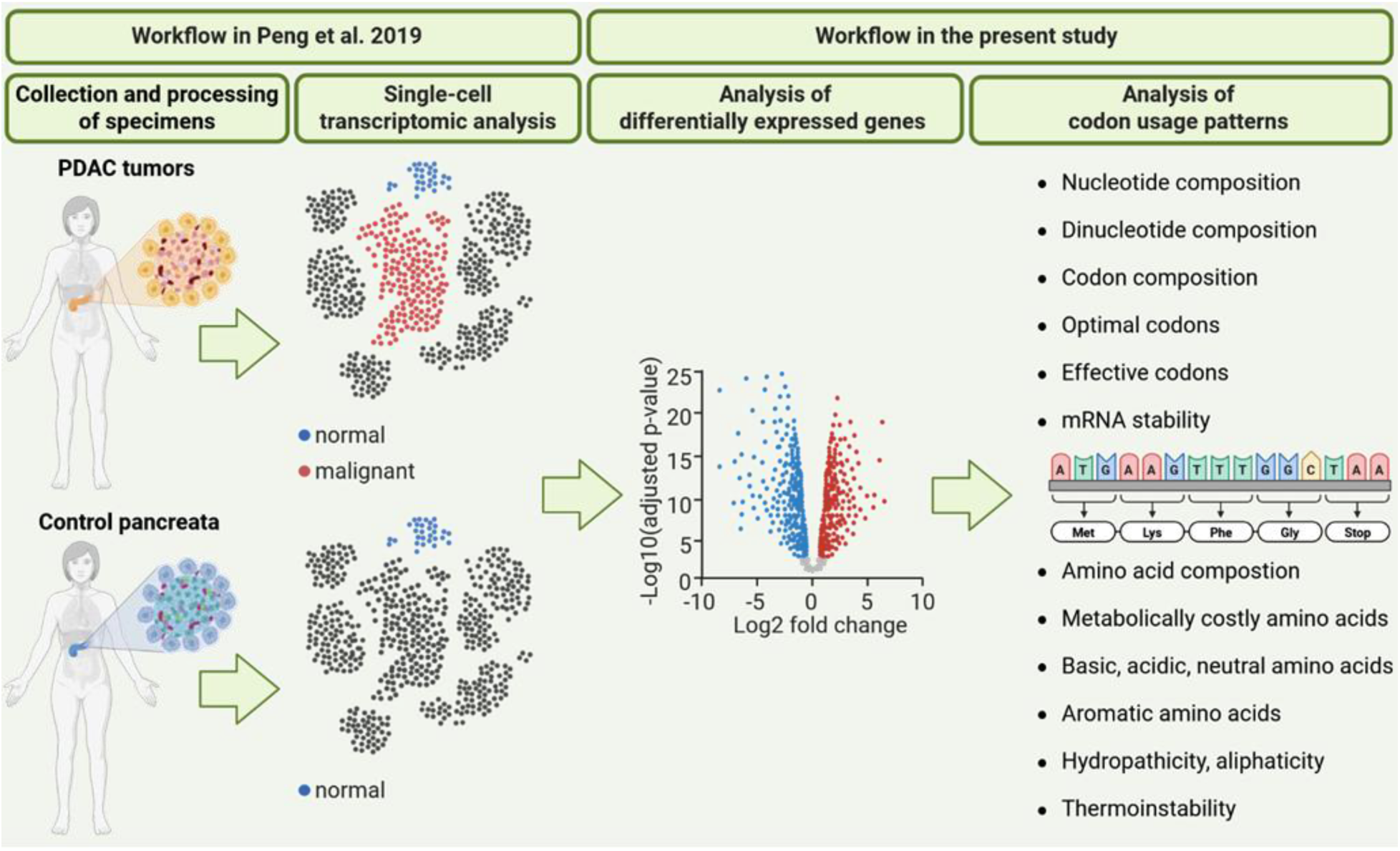
Workflow depicting analyses of codon usage patterns based on single-cell transcriptomics in PDAC.

## RESULTS

To investigate codon usage patterns in malignant PDAC cells, we analyzed publicly available single-cell RNA-seq data (11). In this recent study, Peng *et al.* acquired single-cell transcriptomes from 24 PDAC and 11 healthy pancreatic tissue specimens. Transcriptome-based cell clustering revealed two clusters of ductal cells, termed ductal type 1 and ductal type 2, the latter restricted exclusively to PDAC samples. Differential gene expression analyses performed by the authors reported higher expression levels of genes involved in normal pancreatic ductal functions, such as digestion, pancreatic secretion, and bicarbonate transport (11), in type 1 cells. In contrast, type 2 ductal cells were characterized by upregulation of genes related to proliferation, migration, and hypoxia, suggesting that type 2 ductal cells belong to a malignant subtype. Since ductal type 2 cells showed malignant gene expression profiles and were only detected in PDAC samples (but not in control samples), this cell subtype will henceforth be referred to as ‘malignant’, whereas ductal type 1 cells will be defined as ‘normal’ (**Figure 1**).

Re-analysis of gene expression data from Peng *et al.* (11) identified 2,005 upregulated and 2,947 downregulated DEGs in malignant compared with normal cells (FDR-adjusted p < 0.05, |log2FC| ≥ 1) (**Figure 2A**, **Supplementary Table 1**). After excluding genes lacking sequence information in the ENSEMBL database, 1,774 upregulated and 2,315 downregulated DEGs were analyzed for codon usage bias using their longest coding sequences (CDS).

**Figure 2:**
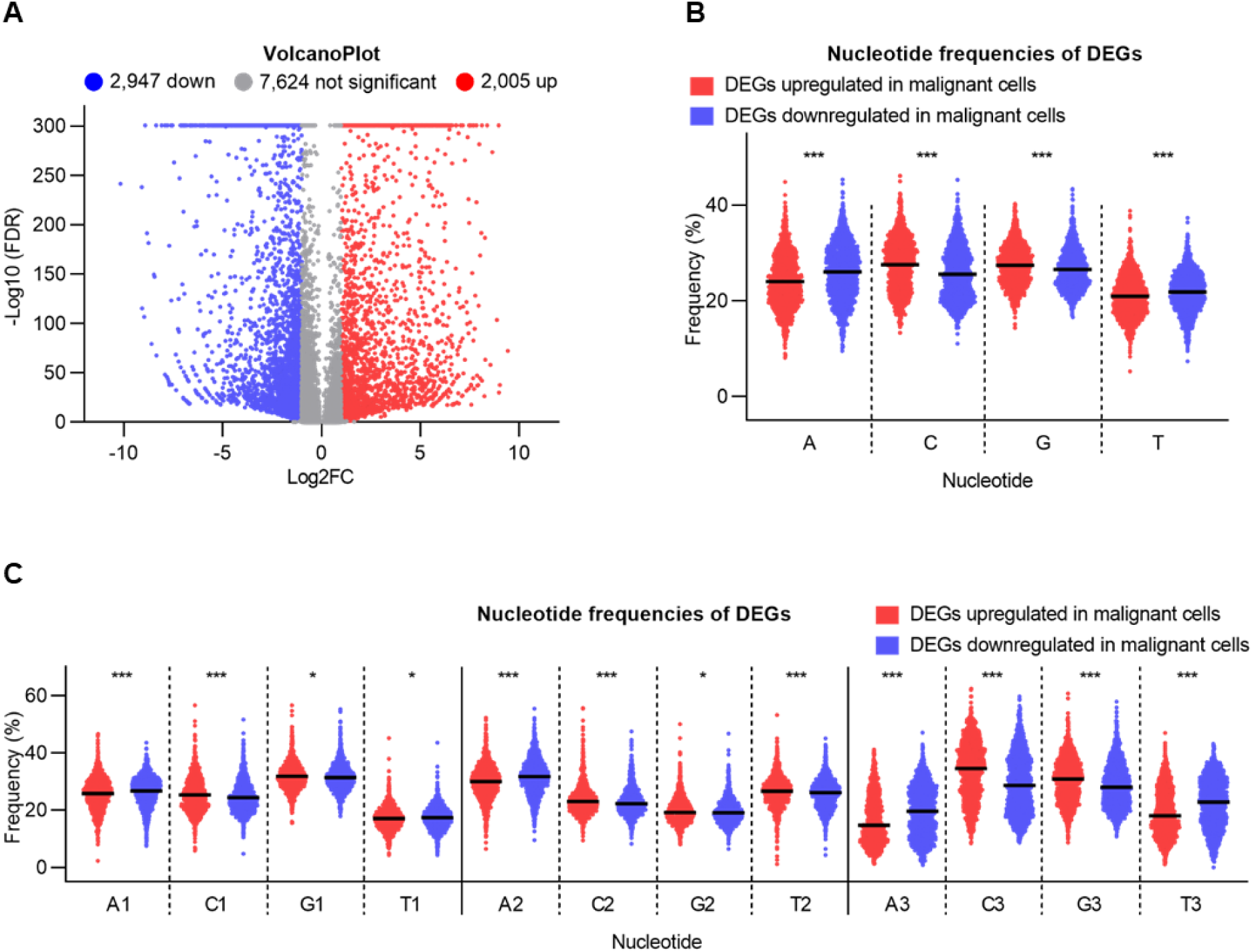
Nucleotide composition bias of differentially expressed genes (DEGs) up- or downregulated in malignant cells. **(A)** Volcano plot showing DEGs. Grey points indicate non-significant changes in expression, while red and blue points represent upregulated and downregulated genes, respectively. **(B)** Percentage of overall base content (A, C, G, T). **(C)** Percentage of nucleotide composition at the first, second, and third codon positions (A1-3, C1-3, G1-3, T1-3).

To identify enriched functional categories among the pre-selected DEGs, we performed an over-representation analysis. Consistent with the findings of Peng *et al.* (11), our analysis revealed that genes upregulated in malignant cells were primarily associated with cell adhesion and proliferation, whereas downregulated genes were associated with metabolism and pancreatic secretion (**Supplementary Figure S1A**). The enrichment of pancreatic-related cell type identifiers (**Supplementary Figure S1B**) and curated gene sets (**Supplementary Figure S1C**) further validated our pre-selection and served as a quality control measure.

Having confirmed the validity of our DEG selection, we next analyzed codon usage patterns based on the longest CDS of up- and downregulated DEGs to characterize malignant PDAC development.

### Nucleotide composition

Since codon usage patterns are influenced by several factors, including compositional bias in nucleotide sequences, we first calculated the nucleotide base composition for all CDS of DEGs up- and downregulated in malignant cells. In genes upregulated in malignant cells, the bases C and G occurred more frequently than A and T (**Figure 2B**). Conversely, genes downregulated in malignant cells displayed significantly higher A and T frequencies.

Further positional effect analysis of nucleotide composition showed significant AT/GC differences at all three codon positions between DEGs in malignant versus normal cells. However, differences in the third codon position were clearly more pronounced, with C3 or G3 showing the strongest enrichment in upregulated DEGs, while A3 and T3 were more prevalent in downregulated genes (**Figure 2C**). This suggests that overall base composition of the DEGs was strongly influenced by the nucleotides occurring at the third codon position.

Collectively, these results reveal a strong bias towards G and C bases in genes upregulated in malignant cells.

### Dinucleotide composition

Because certain dinucleotides are known to exhibit compositional bias in cellular genomes (14, 15), we further quantified dinucleotide representation in the CDS. Under random nucleotide usage, the frequencies of all 16 dinucleotide pairs should be approximately equal. Consistent with the higher overall G and C nucleotide content (**Figure 2B**), C- and G-containing dinucleotides also occurred more frequently than A- and T-containing dinucleotides in genes upregulated in malignant cells (**Figure 3A**).

**Figure 3:**
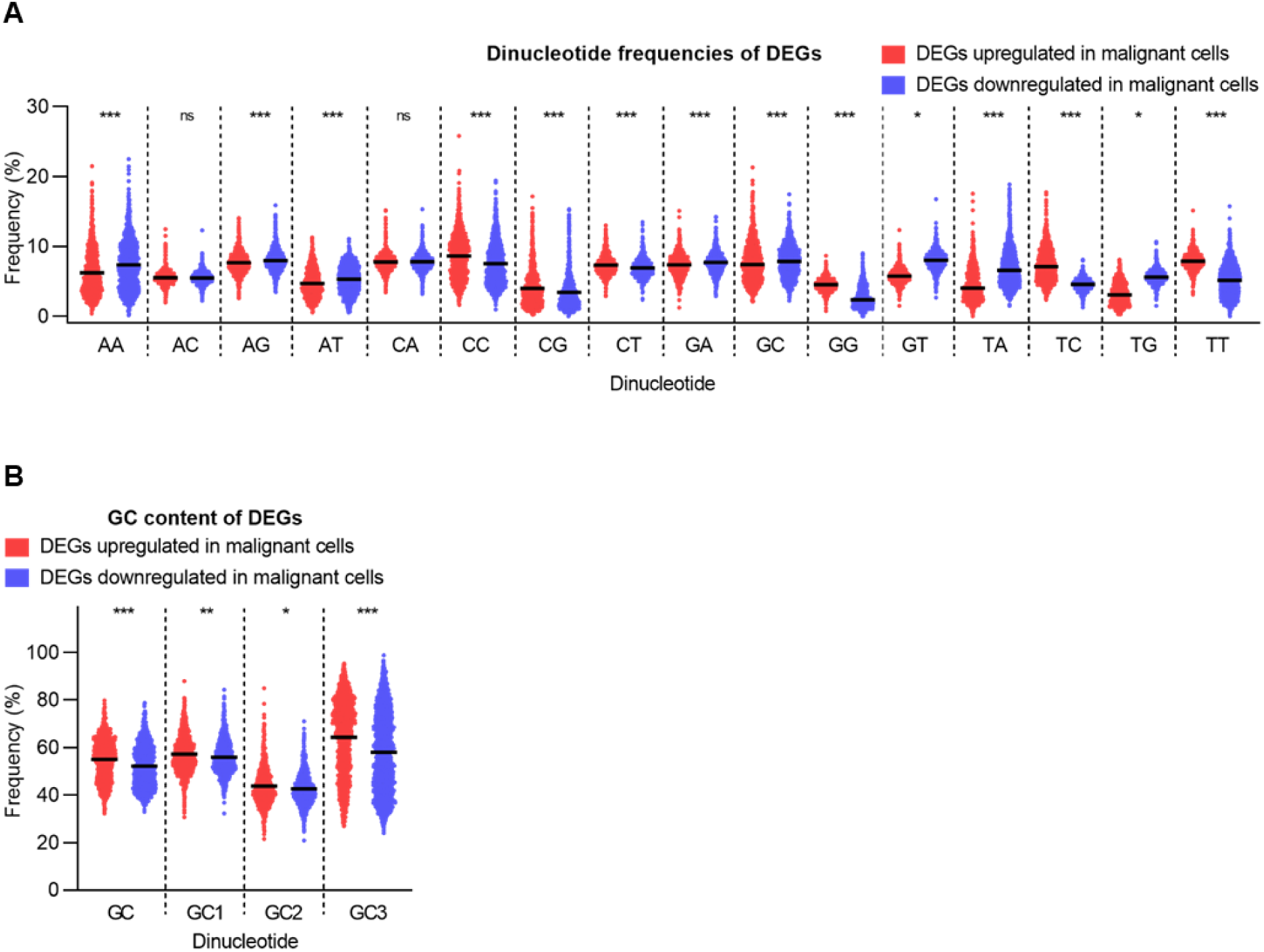
Dinucleotide composition bias of differentially expressed genes (DEGs) up- or downregulated in malignant cells. (A) Distribution of dinucleotide content. (B) Overall GC content and percentage of GC composition at all three codon positions (GC1-3).

GC composition has been reported as a significant factor influencing codon usage bias across genomes (16). Our data revealed that genes upregulated in malignant cells were enriched in GC content, as indicated by a significant increase in overall GC and GC3 content (**Figure 3B**).

In summary, these results reveal distinct patterns in dinucleotide usage, characterized by an enrichment of GC content, particularly at the third codon position, in the genes upregulated in malignant cells.

### Codon composition

Gene expression can be strongly influenced by codon composition. To examine heterogeneous codon usage patterns efficiently, we evaluated the relative synonymous codon usage (RSCU), which is a key index that compares the observed frequency to its predicted frequency within the synonymous codon family coding for a specific amino acid (17). RSCU values greater than 1.6 and less than 0.6 indicate overrepresented and underrepresented codons, respectively, whereas randomly used codons are indicated by RSCU values between 0.6 and 1.6 (17). The results revealed prevalent usage of three codons (CTG, GTG, GCC) in genes upregulated in malignant cells, whereas 11 codons (CGA, ACG, CCG, GCG, CAA, CGT, TCG, ATA, CTA, GTA, TTA) were underrepresented in these genes (**Figure 4A**). In contrast, in genes downregulated in malignant cells, only two codons (CTG, GTG) were overrepresented, while ten codons (CAA, GTA, TTA, ATA, ACG, CCG, CGT, GCG, CTA, TCG) were underrepresented (**Figure 4A**). This substantial overlap in over- and underrepresented codons between both gene sets suggests that RSCU primarily reflects a general human codon usage bias rather than a cancer-specific signature. To overcome this limitation and specifically isolate the differential codon usage between malignant and normal states, we shifted our focus to the relative preference of synonymous codons. Because the genetic code is degenerate and each amino acid – except for methionine (Met, M) and tryptophan (Trp, W) – is encoded by varying numbers of synonymous codons, we analyzed whether certain codons are preferentially used over others for specific amino acids. To perform a differential codon usage analysis, we calculated the odds ratio based on RSCU values for each codon. This approach allowed us to compare their usage frequency directly between upregulated and downregulated DEGs in malignant cells. An odds ratio greater than 1 indicates that a given codon is used at a higher frequency in upregulated compared to downregulated DEGs, whereas an odds ratio below 1 reflects preferential usage in downregulated genes. A marked bias towards G- and/or C-ending codons in genes upregulated in malignant cells, and a corresponding preference for A- and/or T-ending codons in downregulated genes (**Figure 4B**), confirmed the GC3 enrichment in upregulated DEGs (**Figure 3B**) and strongly indicated non-random usage of synonymous codons for specific amino acids.

**Figure 4:**
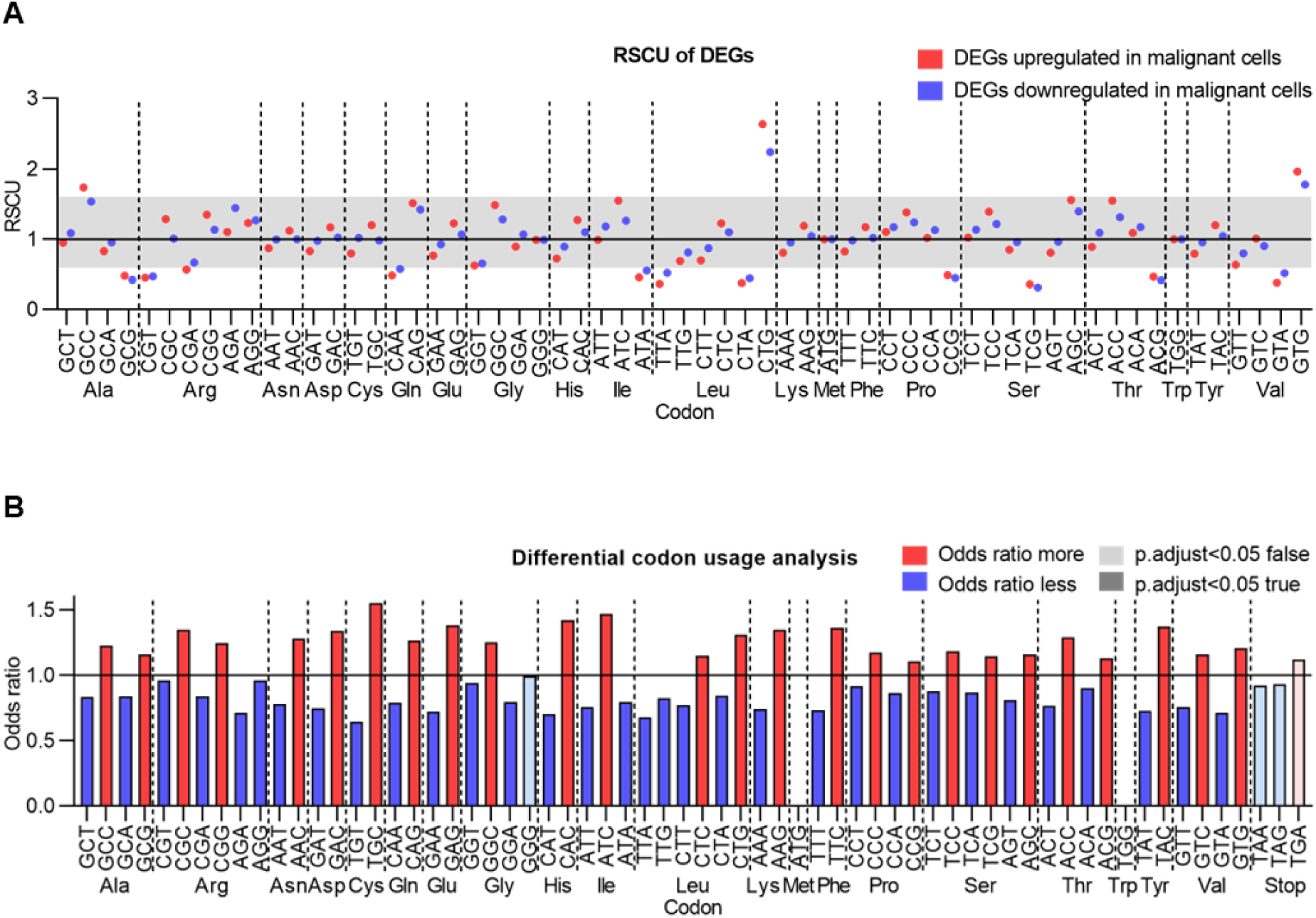
Codon composition bias of differentially expressed genes (DEGs) in malignant cells. **(A)** Scatterplot showing RSCU values for up- and downregulated DEGs. Codons plotted within the white-shaded regions indicate overrepresentation (RSCU > 1.6) or underrepresentation (RSCU < 0.6). **(B)** Differential codon usage analysis based on RSCU calculations; an odds ratio > 1 indicates that a given codon is used at a higher frequency in upregulated compared to downregulated DEGs. Family-wise tests were performed to statistically examine whether synonymous codons are used differentially relative to other codons that encode the same amino acid.

Taken together, these data suggest that the bias observed at the nucleotide and dinucleotide levels is directly reflected at the codon level, where genes upregulated in malignant cells exhibited a distinct preference for G/C-ending codons. This systematic shift in codon usage aligns with the enrichment of G and C bases previously identified in the malignant cell transcriptome.

### Codon usage bias is a main driver of gene expression rates

The preferential or non-random use of synonymous codons plays a central role in optimizing gene expression. To assess the degree of codon usage bias in detail, we calculated two common metrics: the fraction of optimal codons (FOP) and the effective number of codons (ENC).

FOP describes the ratio of optimal codons (the most frequently used synonymous codon for each amino acid) to the total number of codons (18). The score ranges from 0 (non-usage of the most frequent codon) to 1 (exclusive usage of the most frequent synonymous codon), whereby higher FOP values reflect a stronger preference for particular codons. While the identities of optimal codons were largely shared between up- and downregulated genes (**Supplementary Figure S2A**), their frequency of usage varied markedly. Genes upregulated in malignant cells exhibited higher FOP values than downregulated genes (**Figure 5A**). These results suggest that a preference for specific ‘optimal’ codons may regulate gene expression rates, aligning with the systematic enrichment of G/C-ending codons observed in upregulated DEGs (**Figure 4A**).

**Figure 5:**
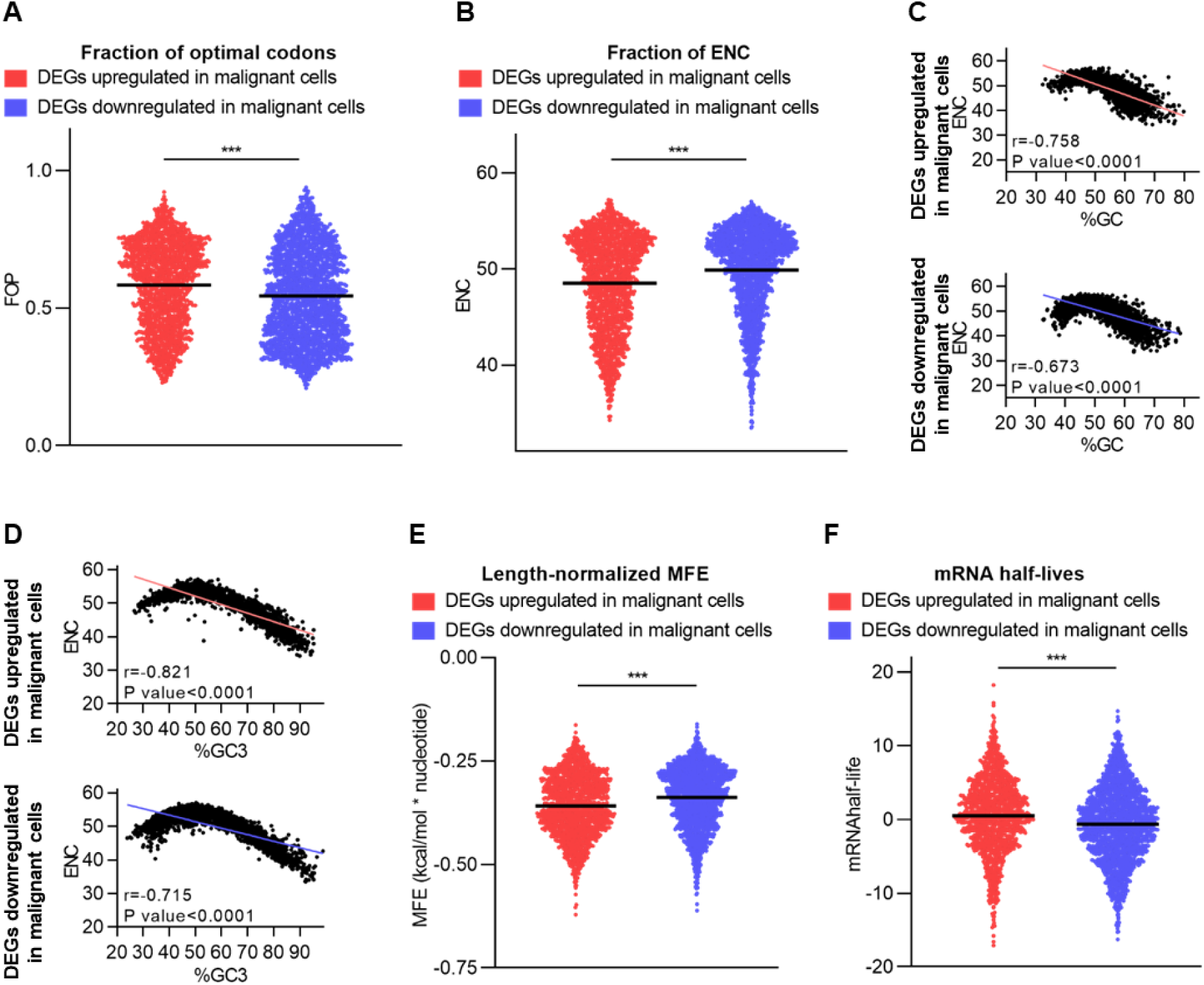
Codon usage bias as a main driver of gene expression regulation. **(A)** Fraction of optimal codons (FOP) and **(B)** effective number of codons (ENC) for differentially expressed genes (DEGs) upregulated and downregulated in malignant cells. **(C)** Correlation between overall GC content and ENC. **(D)** Correlation between GC composition at the third codon position (%GC3) and ENC. **(E)** Length-normalized MFE for upregulated and downregulated DEGs. **(F)** Experimentally measured mRNA half-lives derived from a comprehensive meta-analysis of mammalian mRNA decay studies (23).

To quantify the extent of synonymous codon preference, the ENC was calculated. ENC values range from 20 to 61, where a value of 20 indicates maximum bias, with only one codon used per amino acid, whereas a value of 61 indicates random usage of all synonymous codons (19). Accordingly, lower ENC values indicate a stronger codon usage bias. Our analysis showed that upregulated genes displayed a considerably stronger codon usage bias than downregulated genes as indicated by reduced ENC values (**Figure 5B**). To further investigate the drivers of this synonymous codon usage, ENC values were plotted against overall GC content (**Figure 5C**) and GC3 values (**Figure 5D**), revealing marked negative correlations in both up- and downregulated DEGs. However, this correlation was more pronounced in upregulated than in downregulated genes. These data imply that GC content, particularly at the third codon position, contributes substantially to the codon usage bias observed in the malignant cell transcriptome.

Beyond nucleotide composition, mRNA degradation rate is a major determinant of protein levels in gene expression, and mRNA stability can be strongly influenced by codon usage bias: stable mRNAs are enriched in optimal codons, whereas unstable mRNA molecules typically contain non-optimal codons that promote destabilization (20). Since optimal codons were enriched in genes upregulated in malignant cells (**Figure 5A**), we hypothesized that these genes might encode more stable mRNAs – although enhanced translational efficiency cannot be excluded as a contributing mechanism.

The estimated mRNA minimum free energy (MFE) – the lowest thermodynamic free energy an mRNA sequence can attain when folded into its most stable secondary structure – is a crucial factor in this context. Lower MFE values reflect more stable secondary structures with higher resistance to degradation, ultimately leading to prolonged mRNA half-lives (21, 22). Genes upregulated in malignant cells showed significantly lower length-normalized MFE than downregulated DEGs, suggesting enhanced transcript stability (**Figure 5E**). Additionally, correlation analysis between MFE and ENC revealed a highly significant positive correlation, indicating that mRNA stability is closely associated with the strength of codon usage bias (**Supplementary Figure S2B**).

To experimentally validate our computational predictions of enhanced mRNA stability in genes upregulated in malignant cells, we analyzed mRNA half-lives from a comprehensive meta-analysis of mammalian mRNA decay studies (23). Genes upregulated in malignant ductal cells exhibited significantly longer mRNA half-lives than downregulated genes (**Figure 5F**), confirming our MFE-based stability predictions with experimental data.

In conclusion, the transcriptome of malignant cells is strongly influenced by codon usage bias, characterized by an increased preference for optimal codons and enhanced mRNA stability in upregulated DEGs.

### Amino acid composition

Higher mRNA stability has been linked to increased elongation rates during mRNA translation (24), where more efficient translation increases the amount of protein produced. To analyze whether there is a codon-dependent effect on protein characteristics, various protein-related indices were calculated.

First, amino acid usage analysis revealed that Leu, Ser, Ala and Glu were the most frequent (>7%) in both up- and downregulated DEGs, respectively, while His, Cys, Met, and Trp were the least common (**Figure 6A**). On average, twelve amino acids differed significantly between the two gene sets: Ala, Gly, Leu, Pro, Thr, Trp, and Val were enriched in upregulated DEGs, whereas Arg, Gln, Glu, His, and Lys were enriched in downregulated DEGs.

**Figure 6:**
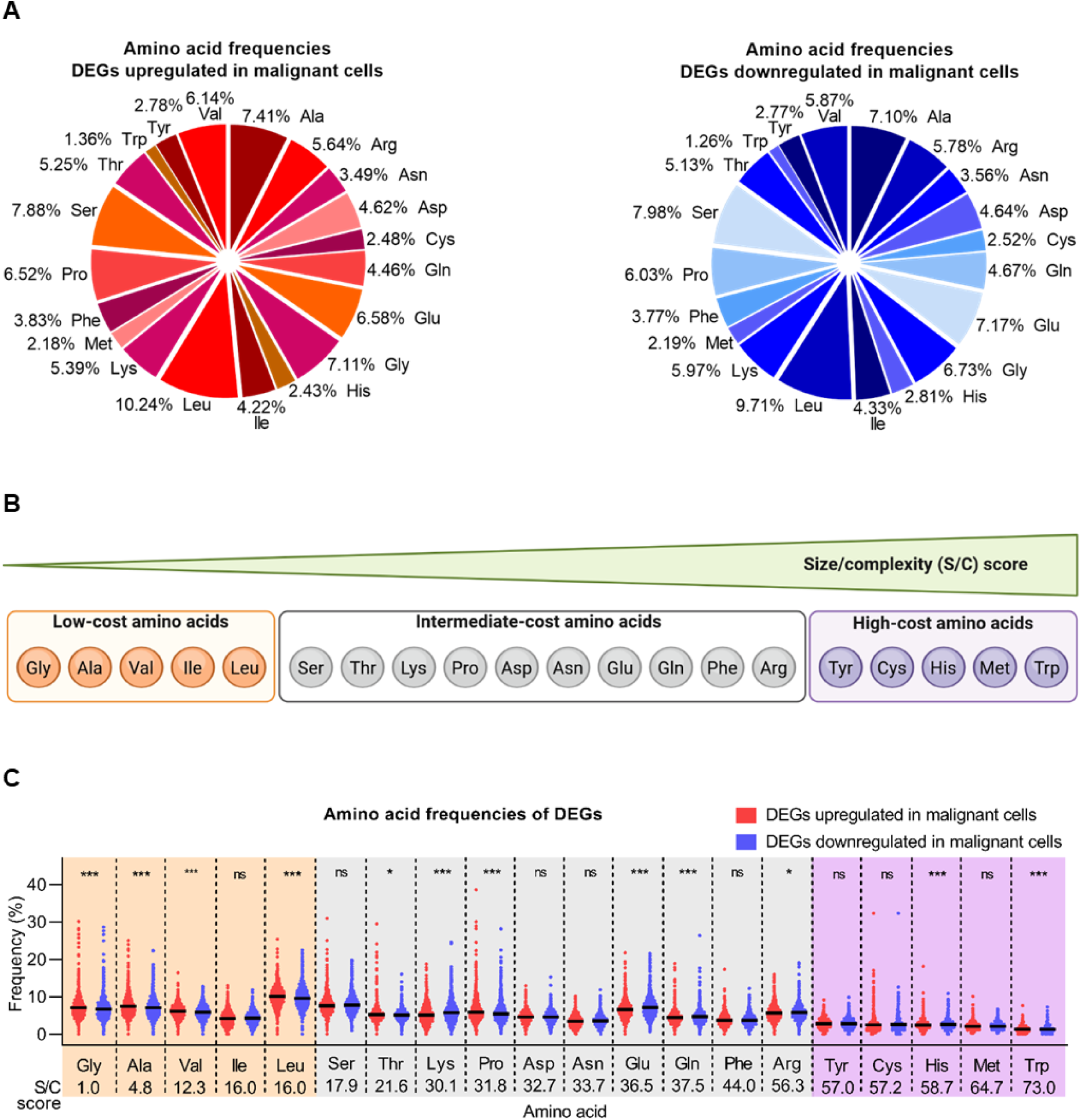
Amino acid composition and metabolic cost bias in malignant versus normal cells. **(A)** Amino acid frequency distribution in genes upregulated (left, red) and downregulated (right, blue) in malignant ductal cells. Pie charts show the relative frequency of each amino acid as a percentage of total amino acid content in the respective gene sets. **(B)** Schematic of amino acids’ metabolic cost stratification based on size/complexity (S/C) biosynthetic cost scores (26). Amino acids are categorized into low-cost (S/C ≤ 16.04: Gly, Ala, Val, Ile, Leu; orange), intermediate-cost (S/C 17.86–56.34: Ser, Thr, Lys, Pro, Asp, Asn, Glu, Gln, Phe, Arg; gray), and high-cost (S/C ≥ 57: Tyr, Cys, His, Met, Trp; purple) groups. **(C)** Percentage of amino acids and associated size/complexity (S/C) scores. The five lowest-cost amino acids are marked in orange, and the five highest-cost amino acids in purple.

Amino acids in highly expressed genes tend to be less metabolically costly, thereby reducing the energy expenditure for protein synthesis (25). To test this, we examined relative biosynthesis costs using size/complexity (S/C) scores, which range from 1.0 (low-cost Gly) to 73.0 (high-cost Trp) based on molecular weight and physicochemical properties (26) (**Figure 6B**). Most low-cost amino acids (S/C ≤ 16.0: Ala, Gly, Ile, Leu, Val) were enriched in upregulated DEGs, whereas high-cost amino acids (S/C ≥ 57.0: Cys, His, Met, Trp, Tyr) were more prevalent in downregulated DEGs (**Figure 6C**), indicating that expensive amino acids are less frequently used in genes upregulated in malignant cells.

Given the described link between amino acid composition and protein structure and function (27, 28), we next analyzed amino acid distributions. Upregulated DEGs showed significantly fewer basic (Lys, Arg, His) and acidic (Asp, Glu) amino acids compared to downregulated genes, while aromatic (Trp, Tyr, Phe) amino acids were unchanged (**Supplementary Figure S3A, B**).

The hydropathicity index (Kyte-Doolittle scale; (29)) was negative in both gene sets but moderately increased in upregulated DEGs, likely reflecting a reduction in polar amino acids, including Ser and Gln (**Supplementary Figure S3C**). The aliphatic index – the relative volume occupied by aliphatic side chains (Ala, Val, Ile, Leu) (30)) – was also moderately increased in upregulated genes (**Supplementary Figure S3D**). Despite this, instability index values (∼48-49) suggested that both gene sets predominantly encode unstable proteins (**Supplementary Figure S3E**).

Collectively, these results reveal that genes upregulated in malignant cells favor neutral, aliphatic, and metabolically less costly amino acids. This metabolic cost stratification suggests that malignant cells optimize their proteome composition to minimize the biosynthetic burden, thereby preserving ATP and biosynthetic capacity for rapid proliferation.

### Analysis of transcript expression in PDAC

Our analyses using the single-cell RNA-seq dataset from Peng *et al*. revealed an increased usage of optimal codons in genes upregulated in malignant cells. To investigate whether codon optimality is further reflected by the usage of distinct transcript isoforms in PDAC tumors, we analyzed transcript-level bulk RNA-seq data from PDAC samples provided by The Cancer Genome Atlas (TCGA) consortium (31). Although bulk RNA-seq data represent a mixture of distinct cell populations and tumor cell content varies between samples, we speculated that differences in transcript usage would still reflect differential transcript regulation within tumor cells.

For each gene, we identified the highest and lowest expressed transcript isoforms based on their median expression across patient samples. To ensure that only genuinely expressed isoforms were included, transcripts with a median TPM of zero were excluded prior to analysis. Focusing on genes with substantial expression differences (≥100-fold), we obtained a set of 3,364 genes for in-depth analysis of sequence composition and structural features.

Although nucleotide frequencies showed minimal differences between transcript isoforms with high and low expression (**Supplementary Figure S4A**), FOP and ENC suggested codon usage patterns similar to those observed in DEGs in malignant cells, albeit with smaller effect sizes (**Supplementary Figure S4B, C**). FOP was significantly increased in highly expressed transcripts compared to their lowly expressed counterparts, consistent with our observation in single-cell data, where genes upregulated in malignant ductal cells exhibited a higher FOP. Likewise, ENC - as a measure of codon usage bias - was reduced in highly expressed isoforms, mirroring the lower ENC found in genes upregulated in malignant ductal cells. Finally, similar to genes upregulated in malignant cells, length-normalized MFE was decreased in highly expressed transcripts (**Supplementary Figure S4D**), indicating enhanced mRNA stability.

In summary, these findings demonstrate that the codon usage-dependent mechanisms observed at the single-cell level are consistently maintained across transcript isoforms in PDAC tumors. The convergence of increased codon optimality (higher FOP, lower ENC) and enhanced thermodynamic stability (lower MFE) across both datasets strongly suggests that PDAC cells leverage synonymous codon choice to systematically stabilize their malignant transcriptome. This multi-level coordination likely ensures high expression of critical oncogenic factors, providing a robust post-transcriptional foundation for tumor progression.

## DISCUSSION

Since its elucidation in the 1960s, the genetic code has been recognized as the molecular blueprint underlying all biological life. It exhibits degeneracy, whereby multiple synonymous codons can encode the same amino acid. While tryptophan and methionine are each specified by a single codon, the remaining 18 of 20 proteinogenic amino acids are encoded by two to six codons. However, these synonymous codons are not utilized randomly; rather, certain codons are consistently preferred over others for specifying a particular amino acid. This universal phenomenon, known as codon usage bias, has emerged as a direct consequence of the genetic code’s degeneracy and represents a fundamental feature of gene expression regulation across all domains of life (32).

The implications of codon usage bias extend beyond mere translational efficiency, as codon usage patterns interact with multiple cellular and molecular factors to modulate gene expression. Cell state (33, 34), promoter architecture (35), tissue specificity (36), and intron presence (37) all contribute to the complex interplay between codon bias and expression levels. Despite the well-established role of codon usage bias in various biological processes (38), its contribution to disease pathogenesis remains largely unexplored. In particular, the relationship between altered codon usage patterns and cancer development has received remarkably little attention, leaving a major gap in our understanding of translational regulation in malignancy.

Recent advances in single-cell transcriptomics have provided unprecedented insights into the cellular heterogeneity of PDAC. Peng *et al*. (11) identified ten main cell clusters in PDAC samples, including two distinct ductal cell subtypes characterized by normal (type 1) and malignant (type 2) gene expression profiles. Notably, control samples contained six main cell clusters, including type 1 but lacking the malignant type 2 ductal cells, highlighting the disease-specific emergence of this population.

Given the potential role of codon usage patterns in modulating the transcriptional and translational landscape of malignant cells, we investigated codon usage bias in genes differentially expressed in malignant ductal cells.

### Factors influencing codon usage bias in pancreatic malignancy

To elucidate the forces shaping codon usage bias in malignant ductal cells, we first examined the fundamental factors known to influence codon usage patterns. Accumulating evidence suggests that codon usage bias is driven by multiple interacting determinants, including genome composition, gene length and expression level, GC content, and mRNA stability (39–41). Among these determinants, GC content warrants particular attention, as it profoundly influences DNA stability through base stacking interactions and the formation of secondary structures. Elevated GC content has been associated with shorter introns (42) and enhanced levels of gene transcription and recombination (43), suggesting a broader role in shaping the genomic architecture and transcriptional regulation beyond codon selection.

Notably, early analyses of vertebrate genomes revealed that genes encoding abundant cellular housekeeping proteins, such as actin, histone, and tubulin, typically exhibit high GC content at the third codon position (GC3) (44). Intriguingly, cellular oncogenes also display elevated GC3 percentages, hinting at a potential link between GC-rich codon usage and oncogenic transformation (45). Furthermore, the genomic context appears to reinforce these patterns: genes with high GC3 content are predominantly embedded within extended GC-rich genomic regions, whereas those with low GC3 are typically surrounded by AT-rich sequences, suggesting that codon usage bias may reflect broader chromosomal organization and compositional constraints (44, 46).

In the present study, our analyses revealed striking differences in overall base composition and nucleotide content at the third codon position between malignant-associated DEGs and their counterparts. Genes upregulated in malignant cells showed marked enrichment in overall GC and GC3 content. Importantly, these differences do not necessarily reflect an active remodeling of codon usage during malignant transformation, but rather the selective upregulation of transcripts that inherently harbor GC-rich codon compositions. The functional question that follows is whether the translational machinery in malignant cells can efficiently decode the codon spectrum of this altered transcript repertoire. The observed GC and GC3 enrichment in upregulated transcripts may nonetheless carry functional implications, including enhanced translational efficiency through optimized tRNA availability (47, 48), increased mRNA stability through secondary structure formation (49), and association with transcriptionally permissive chromatin environments (50).

The differences in GC content observed in genes upregulated in malignant cells were directly reflected in their codon usage patterns. Analysis of RSCU values revealed a pronounced preference for codons terminating in G or C at the third codon position in the malignant cell transcriptome. This bias toward G/C-ending codons is particularly noteworthy in light of previous findings by Gingold *et al*. (47), who demonstrated that codon preferences in human transcriptomes are functionally stratified: cell-cycle genes preferentially utilize codons ending in A or T, whereas pattern-specification genes favor G/C-ending codons. For most amino acids, distinct codons are preferentially employed depending on the functional context, suggesting that codon selection reflects underlying biological programs rather than stochastic variation (47). The pronounced bias at the third codon position exploits the inherent degeneracy of the genetic code, where this position, commonly referred to as the ‘wobble position’, tolerates nucleotide variation without altering the encoded amino acid due to relaxed base-pairing rules during tRNA - mRNA interaction (48). This positional flexibility allows cells to optimize codon usage for translational efficiency and mRNA stability while maintaining the required protein sequence. The preferential enrichment of G and C nucleotides specifically at the wobble position in genes upregulated in malignant cells suggests that cancer cells may exploit this degeneracy to fine-tune gene expression independently of functional constraints imposed by amino acid requirements. Notably, wobble position optimization has been shown to influence translation elongation rates and ribosome occupancy (49), providing a mechanism by which synonymous codon selection can modulate protein production beyond transcript abundance.

Our analysis further revealed that genes highly expressed in malignant ductal cells exhibit strong enrichment for optimized codons. This observation aligns with previous reports demonstrating that codons with strong usage bias are predominantly found in genes encoding highly expressed proteins (50), indicating that codon optimization serves as a mechanism to enhance translational efficiency. The functional significance of optimal codon usage extends beyond translation rate, as codon optimality has been shown to impact ribosome translocation and connect the processes of translation elongation with mRNA decay (20). Importantly, optimal codon content accounts for the coordinated stability observed in mRNAs encoding proteins with related physiological functions (20), demonstrating that codon optimization operates as a precise fine-tuning mechanism to synchronize both mRNA and protein abundance. These findings underscore that the coding sequence contains information beyond the amino acid sequence itself, embedding regulatory signals that govern mRNA stability. Consistent with this principle, genes upregulated in malignant cells showed significantly higher predicted mRNA stability, suggesting that the preference for GC-rich optimal codons not only enhances translational efficiency but also prolongs transcript half-life, thereby amplifying oncogenic protein production through coordinated transcriptional and post-transcriptional mechanisms.

Beyond the effects on mRNA stability and translational efficiency, analysis of codon usage bias in DEGs revealed additional patterns regarding the metabolic costs of amino acids. While the identity of amino acids used remained largely conserved between up- and downregulated genes, significant differences emerged in the frequency of high-cost versus low-cost amino acids. Specifically, genes upregulated in malignant cells showed a preference for amino acids with low biosynthetic costs. This observation fits into a broader picture of metabolic adaptation during tumorigenesis. Comparisons between normal and tumor tissues suggest that the transition to tumor status enhances metabolic efficiency (51). Tumors typically exhibit increased production of short, low-cost, highly expressed proteins compared to matched normal tissues. This shift may represent a selective advantage, enabling tumor cells to utilize resources more efficiently and minimize metabolic stress despite their frequently dysregulated metabolism and increased protein demand required for rapid growth, especially within the nutrient-depleted microenvironment characteristic of PDAC, where metabolic stress is a constant selective pressure. Thus, the preference for optimal codons in genes upregulated in malignant cells appears to serve a dual purpose: enhancing both the efficiency of protein production through improved mRNA stability and translation, and reducing the metabolic burden through strategic amino acid selection.

### Codon usage bias as a central regulator of pancreatic malignancy

Biased codon usage represents a universal feature across all identified genomes and is increasingly recognized as a regulatory mechanism influencing physiological functions. As discussed above, codon usage constitutes one of several factors that modulate gene expression through multiple mechanisms operating at translational, transcriptional, and post-transcriptional levels.

Importantly, tissue-specific patterns of codon usage have been documented in the human transcriptome (52), highlighting the biological relevance of this phenomenon in cellular differentiation and specialization. However, the pathological role of codon usage bias in disease development remains largely unexplored. Emerging evidence suggests disease-specific alterations in codon usage, with genes commonly implicated in primary immuno-deficiencies and cancer displaying overrepresentation of specific codons, such as the CTG codon (45). Furthermore, a pan-cancer study revealed that optimal codons are associated with cellular differentiation, whereas non-optimal codons correlate with proliferation (7). Despite these initial findings, comprehensive analyses of codon usage patterns specifically in PDAC have been lacking.

Our comprehensive analysis of single-cell and bulk RNA-seq data revealed consistent patterns of codon optimization in PDAC (**Figure 7**). In single-cell data, genes upregulated in malignant cells are characterized by GC enrichment, increased codon optimality (higher FOP, lower ENC), enhanced mRNA stability (lower MFE), and preferential utilization of low-cost amino acids. Remarkably, the analysis of highly expressed transcript isoforms in bulk PDAC samples recapitulated key aspects of this optimization pattern, exhibiting increased codon optimality (higher FOP, lower ENC) and enhanced mRNA stability (lower MFE). This multi-level convergence – from differential gene expression to alternative transcript usage – suggests that codon optimization represents a fundamental mechanism supporting the malignant phenotype in PDAC. These findings provide the first systematic characterization of codon usage bias in PDAC and indicate that malignant transformation involves coordinated optimization of codon usage to maximize metabolic efficiency and protein production, thereby supporting the energetic and biosynthetic demands of rapid tumor growth.

**Figure 7:**
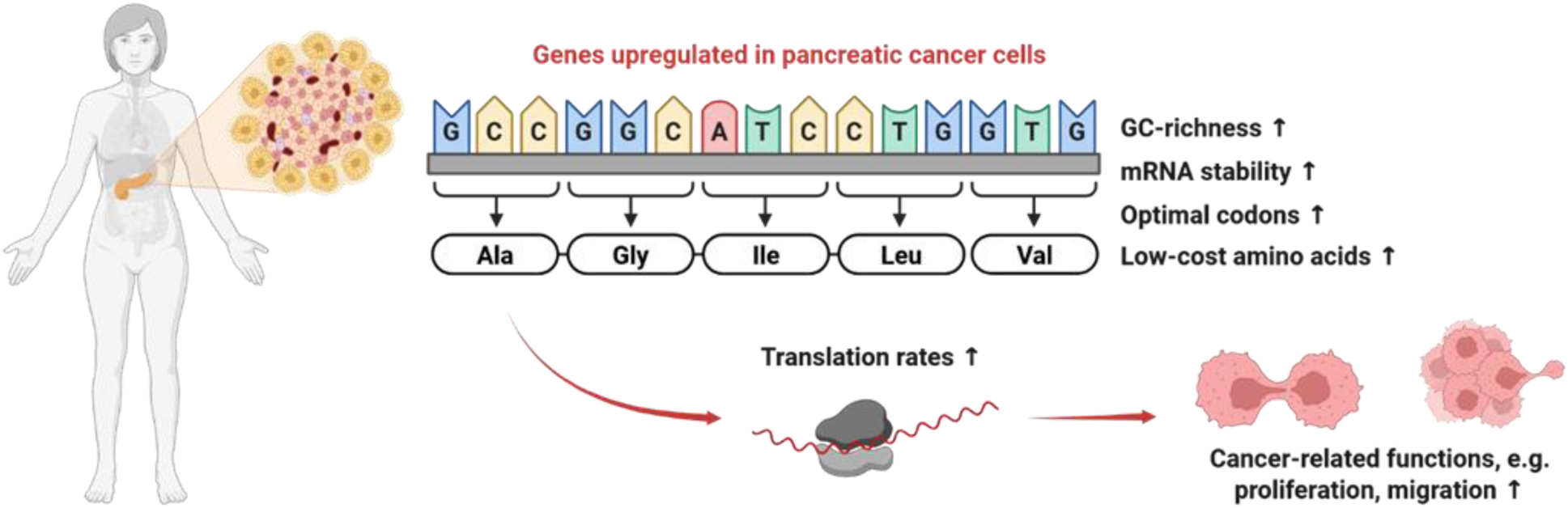
Proposed model depicting the role of codon usage patterns in regulating PDAC malignancy. The schematic illustrates how coordinated optimization of codon usage, mRNA stability, and metabolic efficiency collectively supports the biosynthetic and energetic demands of malignant ductal cells.

## LIMITATIONS AND FUTURE PERSPECTIVES

While our findings provide compelling evidence for codon optimization in PDAC, several limitations warrant consideration. First, our study primarily relies on transcriptomic proxies to infer translational dynamics. Although mRNA stability and codon optimality are tightly coupled, the actual translational output is further modulated by the cellular tRNA pool and its dynamic modifications, the aminoacylation capacity of tRNA synthetases, and the availability of amino acids as substrates for translation. While we observe a bias toward G/C-ending codons, the availability of corresponding tRNAs and the extent of wobble-base modifications could significantly alter decoding efficiency in ways not fully captured by RSCU values alone (3, 53). Furthermore, the relationship between codon optimality and protein fitness is complex. While optimal codons generally enhance elongation rates, non-optimal codons often play a crucial role in programmed translational pausing, which is essential for proper co-translational protein folding (54). Finally, the observed GC-enrichment raises the question of causality versus correlation: it is unclear whether these patterns are primary drivers of oncogenesis or secondary consequences of the altered mutational landscape and genomic instability characteristic of PDAC (55). Future studies integrating Ribo-seq (56) and tRNA-sequencing (57, 58) will be essential to decouple these regulatory layers and confirm the functional impact of codon bias on the PDAC proteome.

## CONCLUSIONS

This study establishes codon usage bias as a critical regulatory layer in PDAC pathogenesis. Our multi-level analysis reveals that malignant transformation systematically exploits synonymous codon degeneracy to optimize gene expression beyond what amino acid sequences alone dictate. The convergent patterns observed across differential gene expression and alternative transcript usage demonstrate that codon optimization is not incidental but represents a fundamental adaptive mechanism in PDAC. By preferentially utilizing GC-rich optimal codons, malignant cells achieve coordinated enhancement of translational efficiency, mRNA stability, and metabolic economy, which collectively support the biosynthetic demands of aggressive tumor growth. Understanding how PDAC cells exploit codon usage bias opens new therapeutic avenues targeting tRNA availability, codon-anticodon pairing, or ribosomal selectivity to selectively impair malignant cell protein production. Such strategies may offer novel approaches for a disease with a persistently poor prognosis and an urgent need for effective treatments.

## METHODS

### Re-analysis of single-cell gene expression data

10X CellRanger output files of human PDAC and healthy control samples, described in Peng *et al*. (11), were obtained from the BioProject website (https://ngdc.cncb.ac.cn/bioproject/browse/PRJCA001063). Functions from the Seurat 5 R-package (59) were utilized for quality filtering, expression level normalization and integration. First, distinct samples were processed independently. Precisely, cells expressing less than 200 distinct genes or whose overall mitochondrial gene expression exceeded 20% (as determined by genes listed in MitoCarta v3 (60)) were removed from further analyses. Genes expressed in less than 20 cells were discarded and potential doublets were detected and removed utilizing the R-package scDblFinder v1.18.0 (61). Expression values were normalized using Seurat’s SCTransform v2 algorithm, employing the 2,000 most variable genes and regressing for mitochondrial gene expression and cell cycle phase. Cell type annotations were assigned according to the information provided by the authors of the data set. Subsequently, samples were integrated by merging individual Seurat objects and performing normalization via SCTransform v2 as described above. Integration was finalized using Seurat’s IntegrateLayers function and the Canonical Correlation Analysis (CCA) method. For differential gene expression analysis, data were prepared using the PrepSCTFindMarkers function, and the identification of DEGs between cell types was conducted applying the FindMarkers function (Wilcoxon Rank Sum test; Bonferroni p-value correction).

### Calculation of nucleotide, amino acid, and codon usage

Transcript sequences were derived by extracting the longest known CDS of the respective genes using the R-packages Biostrings and biomaRt as well as ENSEMBL v110 annotations (62, 63). Nucleotide, amino acid, and dinucleotide frequencies were determined using the Biostrings functions letterFrequency and dinucleotideFrequency. Codon counts, codon differences, ENC, and RSCU were determined using the respective functions of the cubar R-package (64). FOP was calculated using the cubar function get_fop (64), with optimal codons defined separately for each gene set (upregulated and downregulated DEGs) based on the most frequently used synonymous codon within the respective set (op = NULL). This approach does not rely on a shared external reference, which limits the direct comparability of absolute FOP values between groups.

### Calculation of mRNA stability

MFE of each CDS was estimated using RNAfold v2.5.1 (65) and normalized by dividing the resulting MFE value by the sequence length (number of nucleotides). To validate our computational mRNA stability predictions, we utilized experimentally measured mRNA half-life data from a comprehensive meta-analysis of mammalian mRNA decay studies (23). This dataset integrates multiple genome-wide mRNA decay measurements across human cell lines and provides principal component-derived half-life estimates (PC1 scores) that summarize mRNA stability across experimental conditions, excluding outlier datasets. Half-life PC1 scores were mapped to our lists of upregulated and downregulated DEGs in malignant ductal cells by gene symbol, yielding half-life data for 1,390 upregulated and 1,991 downregulated DEGs.

### Over-representation analyses

Over-representation analysis (ORA) was performed using the R-package clusterProfiler (66) and MSigDB gene set collections (v2025.1; (67)). The entirety of gene symbols from the respective gene set collections was used as the background for all enrichment tests.

### Determination of high- and low-abundant PDAC transcripts

Re-analyzed transcript-level Transcripts Per Million (TPM) values of patient samples from the TCGA PAAD cohort were obtained from the study by Tatlow and Piccolo (31). The dataset was filtered by removing non-PDAC samples according to The Cancer Genome Atlas (TCGA) research network (68). Analyses were restricted to transcripts annotated as protein-coding. Median TPM expression across all samples was calculated for each transcript. Transcripts with a median TPM of zero were removed to avoid spurious results from isoforms not expressed in PDAC. For each gene represented by at least two distinct protein-coding transcript isoforms after filtering, we calculated the expression differences between the isoforms with the lowest and highest median expression values. In cases of ties, one isoform was randomly selected to represent the respective lowest or highest expression level. For the calculation of sequence metrics, we selected transcript pairs exhibiting at least a 100-fold difference in their minimum and maximum median expression levels. CDS information for these transcripts were obtained as described above.

### Statistical analysis

Statistical analysis and plot preparation were performed using GraphPad Prism Software (version 8.4.3). All individual data points are displayed in the plots. For comparisons between two independent datasets, a two-tailed Student’s t-test was employed. To compare more than two independent datasets with normal distribution, one-way analysis of variance (ANOVA) followed by a Tukey’s post-hoc test was used. Significance levels are indicated as follows: *p < 0.05; **p < 0.01; ***p < 0.001; ns, not significant.

## DECLARATIONS

### CRediT AUTHORSHIP CONTRIBUTION STATEMENT

**Lisa Müller:** Writing – original draft, Writing – review & editing, Software, Formal analysis, Data curation, Conceptualization.

**Markus Glaß:** Writing – original draft, Writing – review & editing, Software, Formal analysis, Data curation, Conceptualization.

**Pit Preckwinkel:** Writing – review & editing.

**Stefan Hüttelmaier:** Writing – review & editing, Supervision.

**Monika Hämmerle:** Writing – review & editing, Supervision.

**Tony Gutschner:** Writing – original draft, Writing – review & editing, Supervision, Conceptualization.

## ETHICS DECLARATIONS

### Ethics approval and consent to participate

Not applicable.

### Consent for publication

Not applicable.

## ACKNOWLEDGEMENTS

We thank the Wenming Wu lab for providing publicly available single-cell RNA-seq data via the Genome Sequence Archive. Illustrations in Figure 1, Figure 6B, and Figure 7 were created with biorender.com. We acknowledge the use of Claude (claude-sonnet-4-6, Anthropic, San Francisco, CA, USA) for assistance with language editing and grammar refinement.

