## Supplementary Information for "Multi-level Analysis of Codon Usage Patterns Reveals Systematic Optimization of Oncogenic Gene Expression in Pancreatic Cancer"

### Supplementary Figure S1

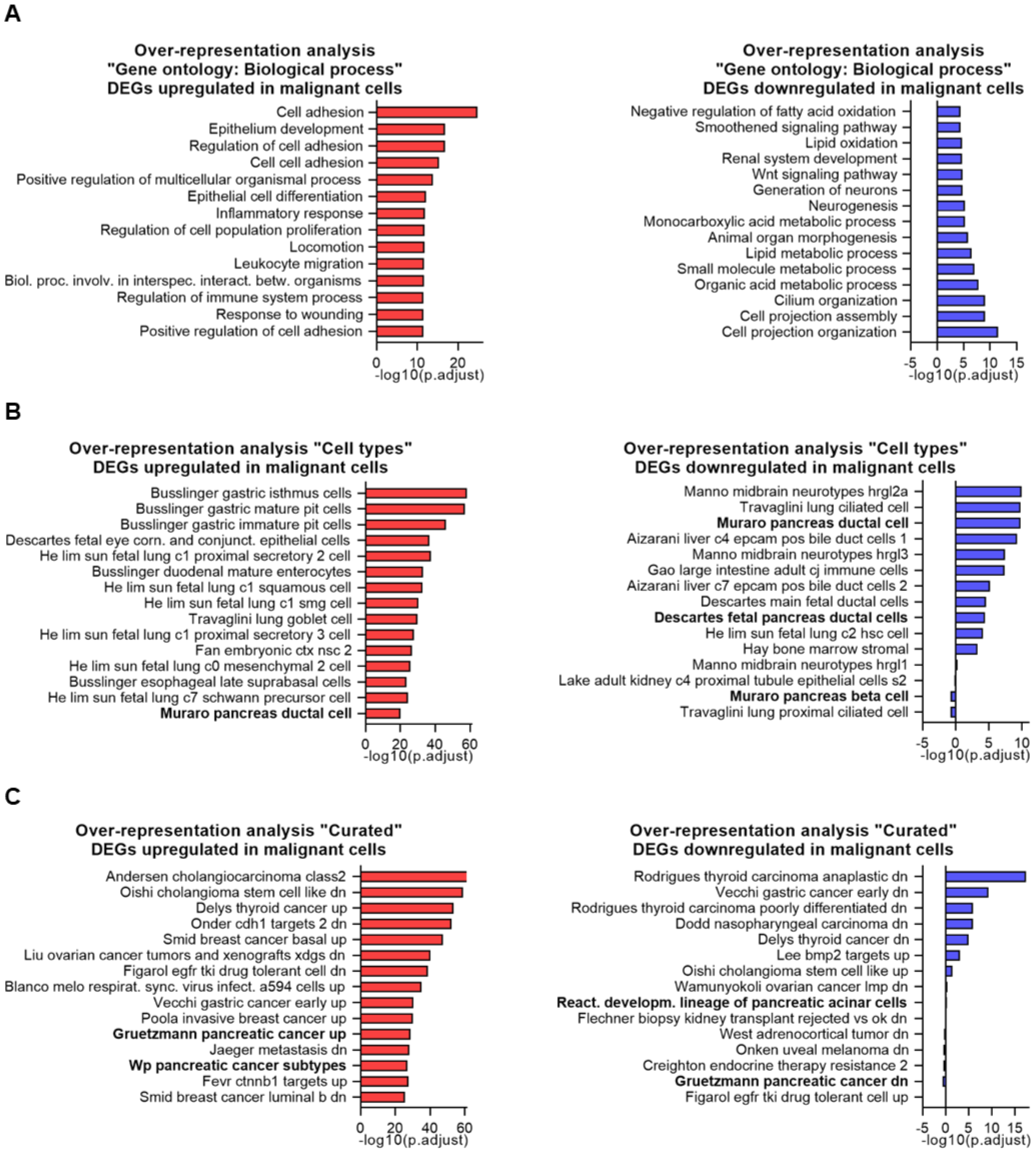


**Supplementary Figure S1:** Over-representation analyses of differentially expressed genes (DEGs) in malignant cells. **(A)** Gene Ontology (GO) Biological Process terms, **(B)** cell type gene sets, and **(C)** curated gene sets. The top 15 enriched gene sets are shown. Pancreas-related terms are highlighted in bold.

### Supplementary Figure S2

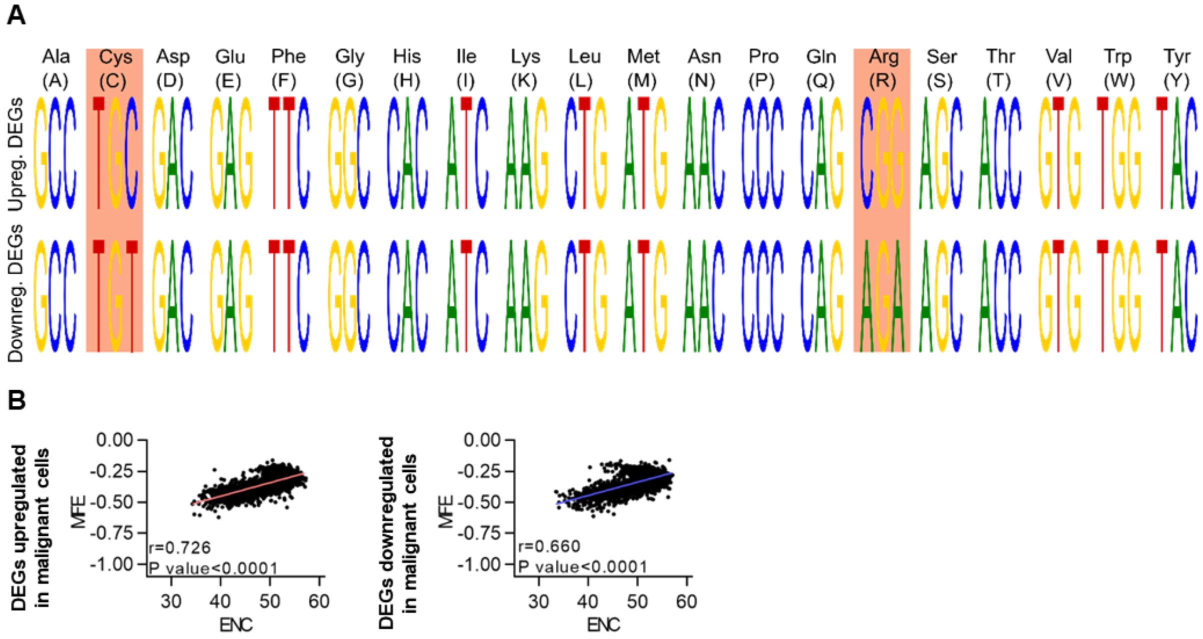


**Supplementary Figure S2:** Bias at the codon level in malignant versus normal cells. **(A)** Graphical representation of codons with the highest RSCU value (defined as optimal codons) for each amino acid among up- and downregulated differentially expressed genes (DEGs) in malignant cells. Differences in codon preference between up- and downregulated genes are marked in orange. **(B)** Correlation between length-normalized MFE and ENC.

### Supplementary Figure S3

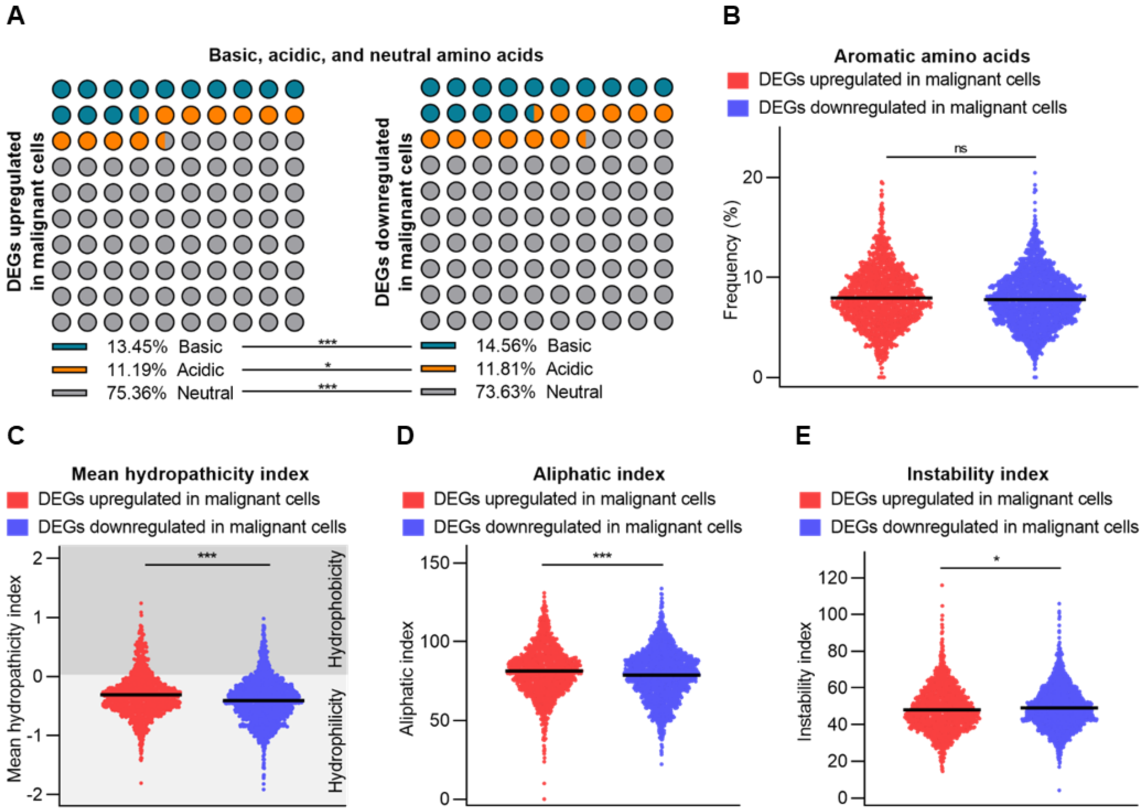


**Supplementary Figure S3:** Amino acid composition bias and physicochemical properties of differentially expressed genes (DEGs) in malignant versus normal cells. **(A)** Percentage of basic, acidic, and neutral amino acids. **(B)** Percentage of aromatic amino acids. **(C)** Mean hydropathicity index based on the Kyte-Doolittle score. **(D)** Aliphatic index. **(E)** Instability index.

### Supplementary Figure S4

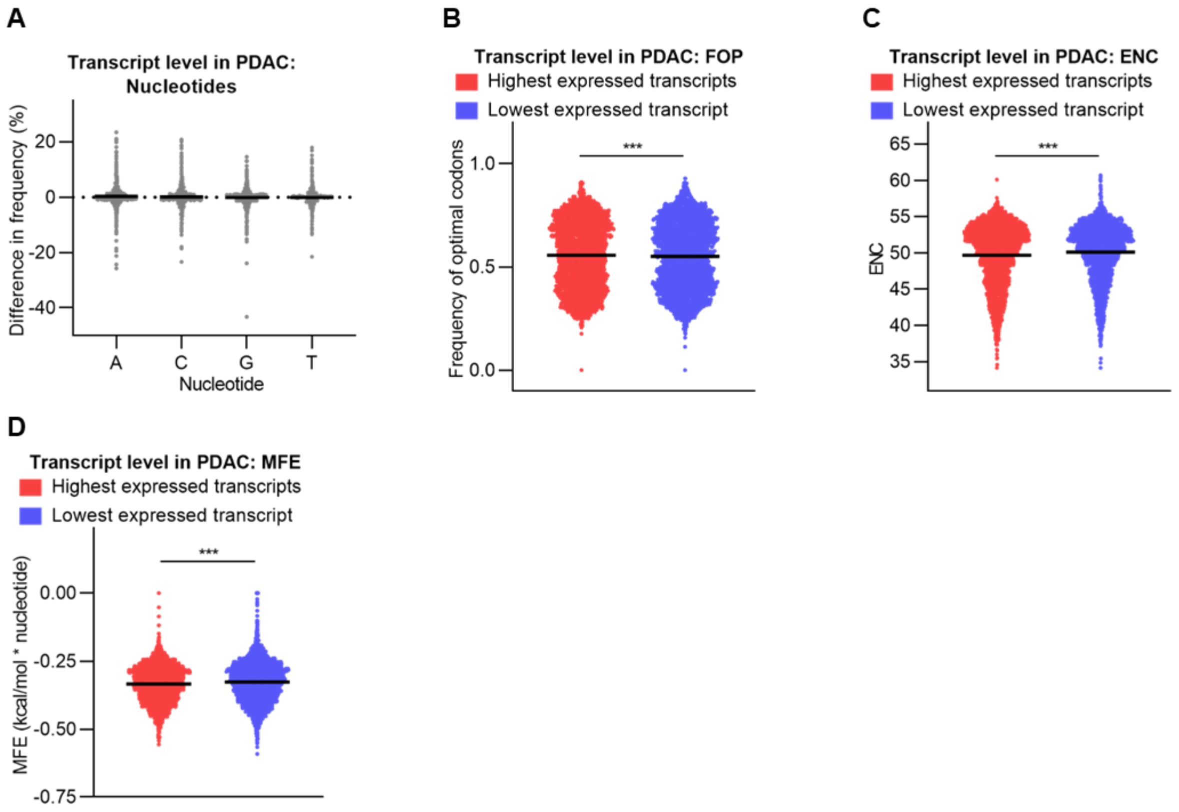


**Supplementary Figure S4:** Codon usage bias metrics and mRNA folding energy in PDAC transcripts with highest and lowest expression levels. **(A)** Differences in nucleotide base frequencies (highest minus lowest expressed transcript isoforms). Positive values indicate enrichment in highly expressed isoforms. **(B)** Fraction of optimal codons (FOP), **(C)** effective number of codons (ENC), and **(D)** length-normalized MFE.
